# Stress- and phospholipid signalling responses in Arabidopsis PLC4-KO and -overexpression lines under salt- and osmotic stress

**DOI:** 10.1101/2023.06.02.543366

**Authors:** Max van Hooren, Essam Darwish, Teun Munnik

**Author notes:** Corresponding author: Teun Munnik, Plant Cell Biology, Swammerdam Institute for Life Sciences, University of Amsterdam, PO Box 1210, 1000 BE Amsterdam, The Netherlands.

## Abstract

Several drought- and salt tolerant phenotypes have been reported when overexpressing (OE) phospholipase C *(PLC*) genes across plant species. In contrast, a negative role for Arabidopsis *PLC4* in salinity stress was recently proposed by Xia et al., (2017) since they reported roots of *PLC4-OE* seedlings to be were more sensitive to NaCl while *plc4*-KO mutants were more tolerant. To investigate this apparent contradiction, and to analyse the phospholipid signalling responses associated with salt stress, we performed root growth- and phospholipid analyses on *plc4*-KO and *PLC4-OE* seedlings subjected to salinity (NaCl) or osmotic (sorbitol) stress, and compared these to wild type (WT) plants. Only very minor differences between PLC4 mutants and WT were observed, which even disappeared after normalization of the data, while in soil, PLC4-OE plants were clearly more drought tolerant than WT plants, as was found earlier when overexpressing Arabidopisis *PLC2, -3, -5, -7* or -*9*. We conclude that PLC4 plays no opposite role in salt-or osmotic stress and rather behaves like the other Arabidopsis PLCs.

**GRAPHICAL ABSTRACT:** 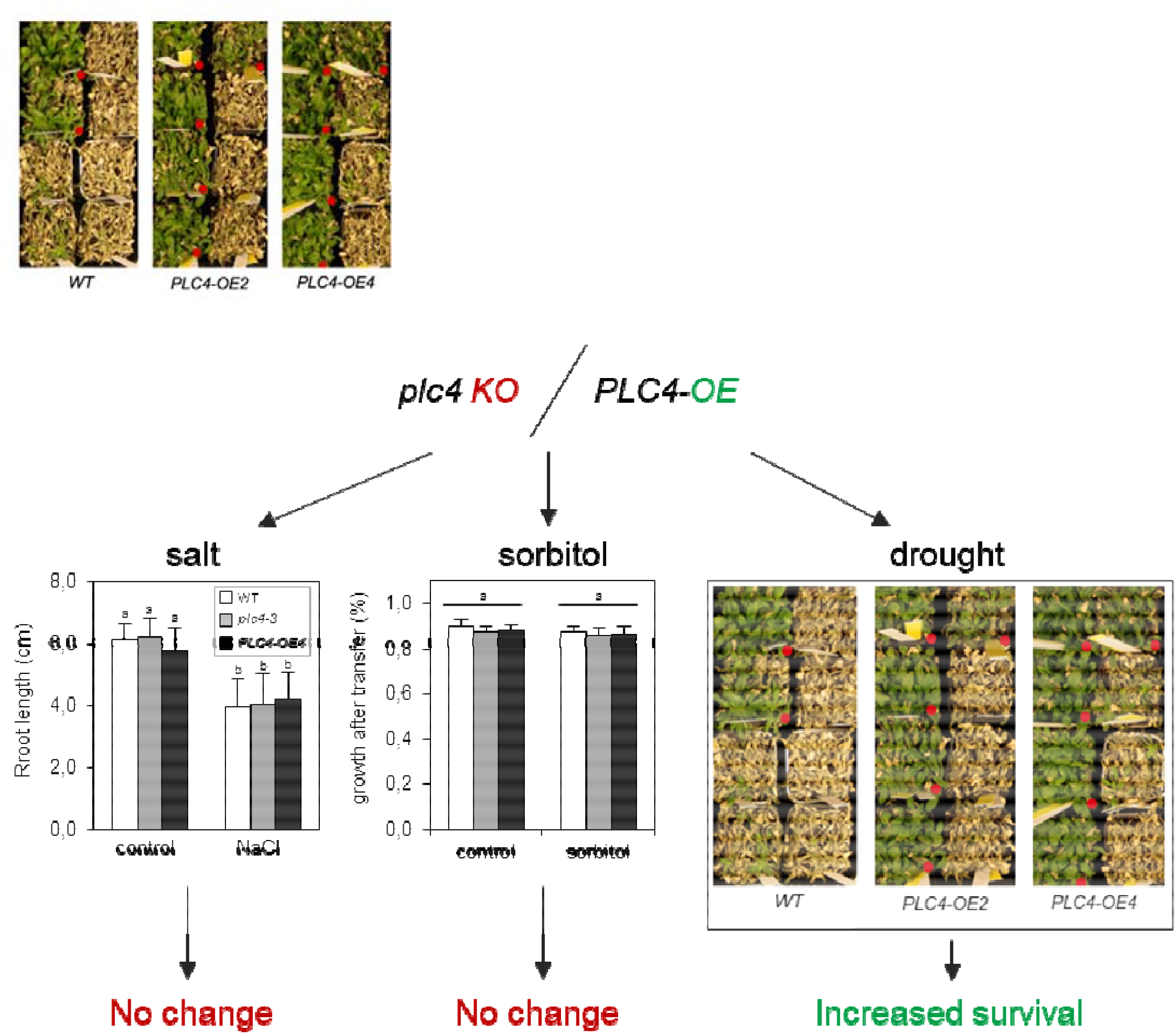

## 1. INTRODUCTION

Plants are sessile organisms and as such have to continuously respond and adapt to their local environment. Understanding how these acclimation reactions take place is of prime importance to secure the world food supply. One of the biggest problems in agriculture is the quality and availability of water (Boretti and Rosa, 2019). Drought and soil salinization are two main problems that are often related as local salt concentrations rise when soils dry out. Hence, pathways involved in drought- and salinity responses often overlap, including the signalling molecules and hormones activating them (Gamalero and Glick, 2022; Golldack et al., 2014; Hao et al., 2022; Marusig and Tombesi, 2020; Verma et al., 2022).

Salt- and osmotic stress typically trigger the formation of two lipid second messengers, i.e. phosphatidylinositol 4,5-bisphosphate (PIP_2_) and phosphatidic acid (PA) (Han and Yang, 2021; Hou et al., 2016; Munnik and Vermeer, 2010; Testerink and Munnik, 2011; Yao and Xue, 2018; Verslues et al., 2023), which are part of the phospholipase C (PLC) pathway. Normally, the concentrations of these lipids are relatively low, in particular of PIP_2_, but *in vivo* ^32^P_i_-radiolabelling experiments revealed that these lipids are rapidly produced in response to salt (NaCl) or osmotic (sorbitol, PEG, mannitol) stress, reacting within seconds to minutes and reaching a maximum at ∼30-60 min (DeWald et al., 2001; Konig et al., 2007; Meijer et al., 2001; Meijer et al., 2017; Munnik et al., 2000; Takahashi et al., 2001; Zonia and Munnik, 2004; van Leeuwen et al., 2007). The formation of both lipids has been associated with important cellular processes, including vesicular trafficking, cytoskeletal reorganization, and transport of molecules across membranes, which are regulated through recruitment and/or activation of specific protein targets, like protein kinases, phosphatases, small G-proteins and membrane transporters (Doumane et al., 2021; Hou et al., 2016; Ischebeck et al., 2013; Lebecq et al., 2022; Naramoto et al., 2009; Noack and Jaillais, 2017; Pleskot et al., 2013; Pokotylo et al., 2014; Pokotylo et al., 2018; Synek et al., 2021; Testerink and Munnik, 2011; Ufer et al., 2017; Wang et al., 2019; Wu et al., 2017; Yao and Xue, 2018; Yperman et al., 2021; Zhao et al., 2010).

For salt- and osmotic stress, a direct interaction between PA and NADPH oxidases, RbohD and RbohF, was uncovered, linking PA with reactive oxygen species (ROS)production (Chapman et al., 2019; Kadota et al., 2015; Zhang et al., 2009). Similarly, PA has been connected to SOS1, an Na^+^/H^+^ antiporter that pumps Na^+^ out of the cell (Qiu et al., 2004). PA binds and activates MKK7 and MKK9, which leads to the phosphorylation and activation of MPK6 (Shen et al., 2019). Alternatively, PA can directly stimulate the phosphorylation, and activation, of SOS1 through MPK6 (Yu et al., 2010).

For PIP_2_, several protein domains that specifically bind its lipid headgroup have been identified, including proteins involved in endo- and exocytosis by interacting with clathrin and EXO70 (de Jong and Munnik, 2021). PIP_2_ formation has been shown to be important for membrane identity and creates cell polarity in the tip of root hairs, pollen tubes and cell plate formation during cell division, having consequences for PIN localization, and the organization of the actin cytoskeleton, in particular at the plasma membrane though PIP_2_ has also been found in the nucleus (de Jong and Munnik, 2021; Dieck et al., 2012; Guo et al., 2020; Ischebeck et al., 2010; Ischebeck et al., 2013; Kato et al., 2019; Mei et al., 2012; Noack and Jaillais, 2020; Song et al., 2021; Wu et al., 2017; Xing et al., 2021; Zhao et al., 2010).

The formation of PIP_2_ is triggered by activation of PIP 5-kinase (PIP5K), which phosphorylates phosphatidylinositol 4-phosphate (PIP) into PIP_2_. For salinity stress, this has recently been shown to involve PIP5K7, -8 and -9 (Kuroda et al., 2021). PLC can hydrolyse both PIP and PIP_2_ as substrates, generating diacylglycerol (DAG) and the water-soluble headgroup, i.e. inositol 1,4 phosphate (IP_2_) or inositol 1,4,5 phosphate (IP_3_), respectively (Munnik, 2014). DAG is rapidly phosphorylated into PA by DAG kinase (DGK) (Arisz and Munnik, 2013), while IP_2_ and IP_3_ can be phosphorylated into higher inositolpolyphosphates (IPPs), like IP_5_ and IP_6_, and even into pyrophosphorylated forms, IP_7_ and IP_8_ (Laha et al., 2016; Munnik, 2014). Abscisic acid (ABA) has been shown to trigger IP_6_ formation in minutes and to release Ca^2+^ from an intracellular store in guard cells, resulting in the closure of stomata (Flores and Smart, 2000; Lemtiri-Chlieh et al., 2000; Lemtiri-Chlieh et al., 2003). A potential role for PLC herein has been confirmed by gene silencing and KO-analyses (Hunt et al., 2003; Mills et al., 2004; van Wijk et al., 2018; Zhang et al., 2018a; Zhang et al., 2018b).

Besides the PLC/DGK route, PA can also be generated through activation of phospholipase D (PLD), which hydrolyses structural phospholipids, like phosphatidylcholine (PC), phosphatidylethanolamine (PE) and phosphatidylglycerol (PG), into PA and the respective alcohol group (choline, ethanolamine or glycerol). Arabidopsis contains 12 PLDs, of which PLDα1 and PLDδ have been implicated in salt- and osmotic stress, as well as in drought (Katagiri et al., 2001; Munnik and Testerink, 2009; Uraji et al., 2012).

Overexpression (OE) of PLC has been shown to improve drought tolerance in various plant species. These include *ZmPLC1* in maize (Wang et al., 2008), *BnPLC2* in canola (Georges et al., 2009), *NtPLCδ1* in tobacco (Tripathy et al., 2012), *OsPLDα1* and *OsPLC4* in rice (Abreu et al., 2018; Deng et al., 2019), *GmPLC7* in soybean (Chen et al., 2021a), and *AtPLC2, AtPLC3, AtPLC5* and *AtPLC7* in Arabidopsis (van Wijk et al., 2018; Zhang et al., 2018a; Zhang et al., 2018b). In rice, overexpression of *PLC* also improved the plant’s tolerance to salt- and osmotic stress (Deng et al., 2019; Li et al., 2017). Similarly, OE of wheat *PLC1* (Ta*PLC1*) in Arabidopsis increased its salt- and osmotic stress tolerance (Wang et al., 2020).

Xia et al. (2017) recently reported that DEX-inducible OE of At*PLC4* in Arabidopsis seedlings resulted in reduced primary root growth under salt stress, while *plc4*-knock-out (KO) mutants showed an improved growth performance under salinity, implying a negative role for PLC4 in salt stress. Considering the above, we found this rather counterintuitive, especially since of all nine Arabidopsis PLCs, *PLC4* has the highest root expression and the highest fold-increase in response to salt stress (Tasma et al., 2008). Hence, we re-analysed the primary root growth of *PLC4-*KO and -OE lines, studied their phospholipid signalling responses when subjected to salt- or osmotic stress and tested the effect of *PLC4*-OE on the drought tolerance of Arabidopsis plants in soil.

## 2. RESULTS

### 2.1. *gPLC4* expression in KO- and OE mutants

To measure the expression level of PLC4 in the various mutant backgrounds, qPCR was used (Fig. 1). *PLC4-OE2* and *PLC4-OE4* displayed 2.7- and 4.2-times higher expression levels than WT, respectively, while the expression in the plc4-3 line, which is the same line as used by Xia *et al*. (2017), was severely reduced and close to zero. Surprisingly, however, *plc4-2*, was found to exhibit *PLC4* levels like WT. Upon further inspection, the T-DNA insertion of *plc4-2* was found to be positioned at an intron, which may explain why there is no reduced expression. For the rest of the experiments, we disregarded this mutant.

**Figure 1.**
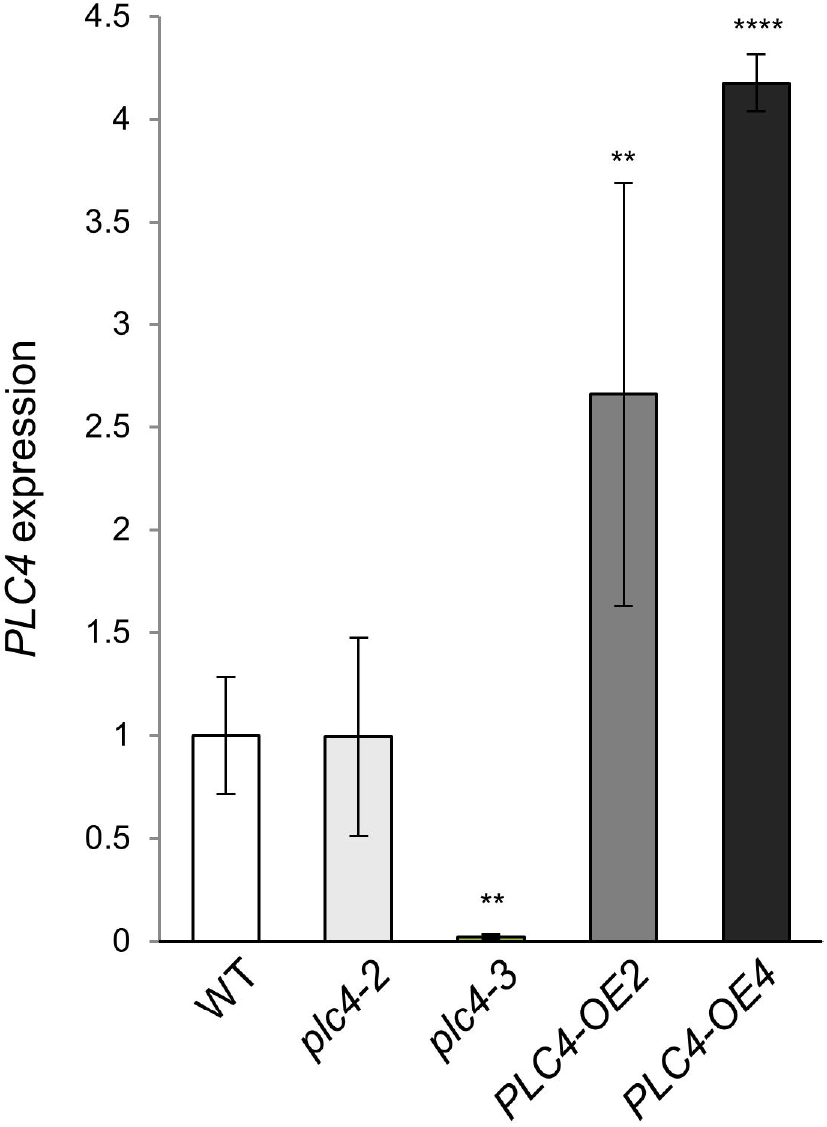
*PLC4* expression in wild-type and *PLC4* mutants. Expression levels measured by qPCR and normalized to SAND. Values are means ± SE of 2 independent experiments with 3 biological replicates each. ANOVA test: **, P<0.01; ****, P<0.0001.

### 2.2. Primary root growth under salt- and osmotic-stress

Next, we compared the primary root length of *PLC4-*KO and -OE lines with WT for their growth under control-, salt- or osmotic stress conditions. Four-day old plants were transferred to agar plates with or without 100 mM NaCl and grown for another eight days (Fig. 2). Plants exposed to salinity stress had significantly smaller roots than control plants (64-72% of control). In contrast to what was found by Xia *et al*. (2017), where *plc4* seedlings grew faster than WT and *PLC4*-OE grew slower, no significant differences between WT, *plc4-3* and *PLC4-OE4* were found. Small, non-significant changes were observed at control conditions, but even then, when normalized, no significant changes were observed (Fig. 2B). Repeated experiments with *PLC4-OE2* instead of *PLC4-OE4* gave similar results, also showing no significant difference in response. As an alternative, we measured how much primary root growth occurred after the transfer to the salt plates, and visualized this as a percentage of total primary root length (Fig. 2D). As such, a small but significant increase of 4% for the *plc4-3* mutant at the salt condition was found. At control conditions, there were small changes too but these were not significant. Normalizing for these small changes, removed all significant changes for the *plc4-3* under salt conditions (Fig. 2E).

**Figure 2.**
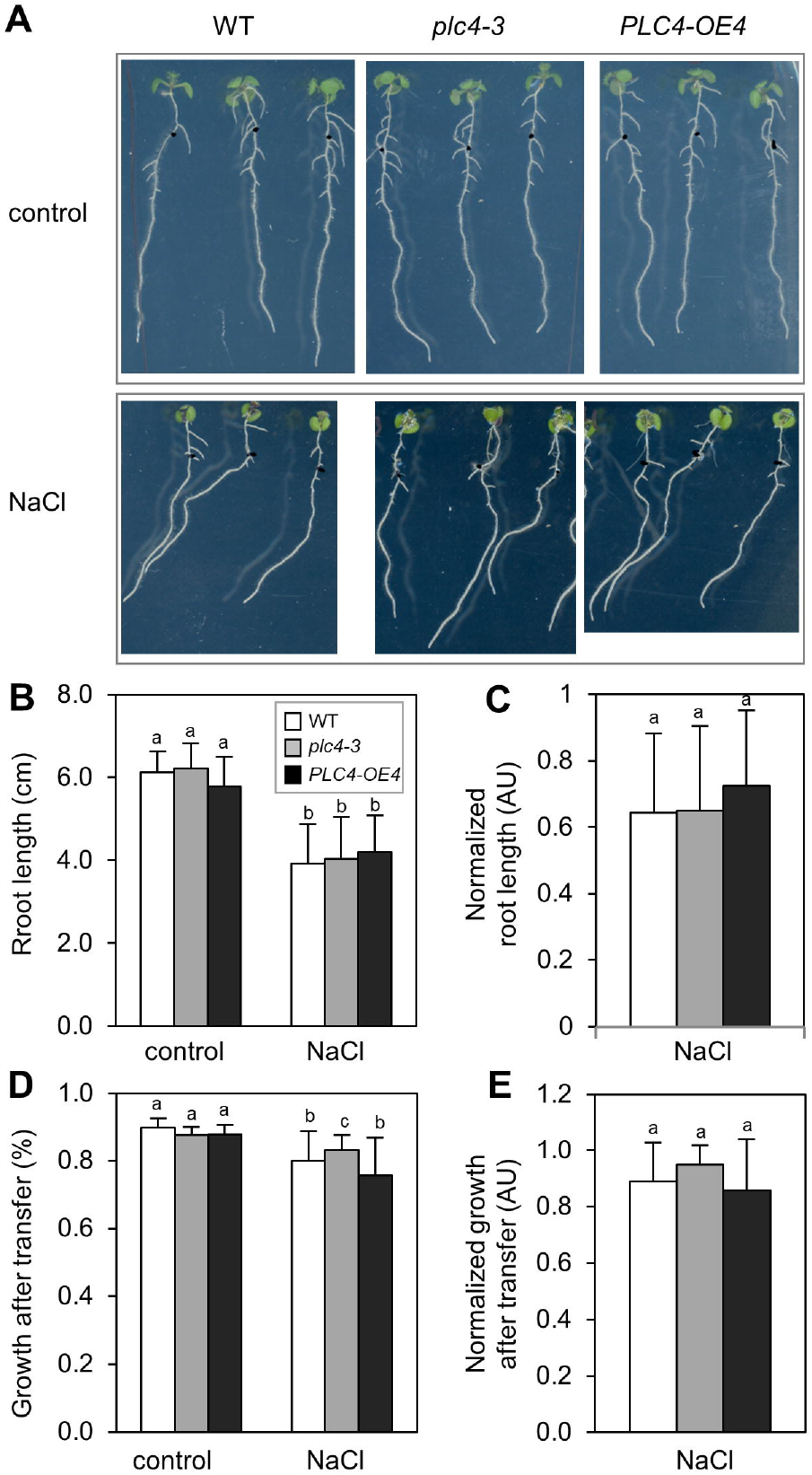
Effect of salt stress on primary root growth of Arabidopsis WT and *PLC4* mutants. Seedlings were grown for 4 days on½MS-agar plates and then transferred to½MS plates supplemented with 100 mM NaCl or control plates without salt. **A)** Seedlings, 8 days after transfer (DAT). **B)** Primary root length 8 DAT (12 days old). **C)** Primary root length normalized to their size at control conditions. **D)** Primary root growth after transfer as ratio of total main root length. **E)** Relative root length normalized to the control conditions in. N=75-99 from 2 independent experiments with error bars representing SE. Statistics were done using two-way ANOVA test and post hoc TUKEY test. Significance group(s) are indicated with letters (P<0.05).

Performing the experiments with 200 mM sorbitol, which is osmotically similar to 100 mM NaCl but without the ionic stress, we found *PLC4-OE4* roots were slightly (4.4%), but significantly, smaller at sorbitol conditions (Fig. 3A). However, the significance was lost when normalizing for their size at control conditions (Fig. 3B). As such, no significant changes were found when measuring primary root-growth responses after transferring (Fig. 3C, 3D).

**Figure 3.**
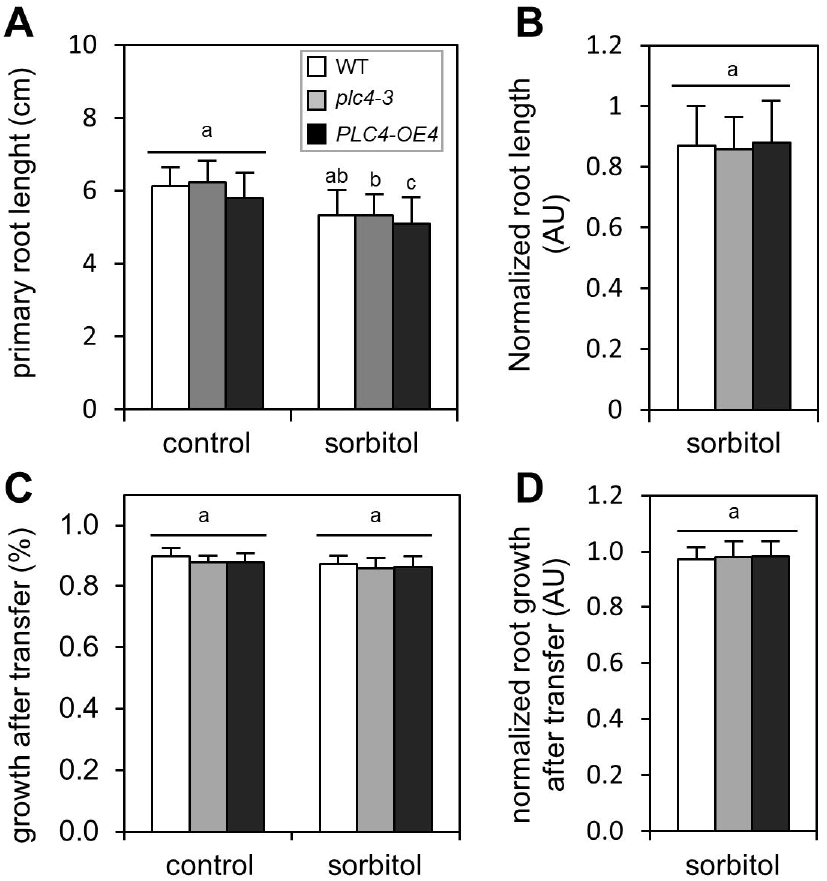
Effect of hyperosmotic stress on primary root growth of WT and *PLC4* mutants. Seedlings were grown for 4 days on½MS plates and then transferred to½MS ± 200mM sorbitol, after which primary root length was measured 8 DAT. **A)** Primary root length normalized to their size at control conditions. **B)** Primary root length normalized to size of genotypes at control conditions. **C)** Primary root growth after transfer as ratio of total main root length. **D)** Relative root length normalized to the control conditions in. N=75-99 from 2 independent experiments with error bars representing SE. Statistics were done using two-way ANOVA test and post hoc TUKEY test. Significance group(s) are indicated with letters (P<0.05).

### 2.3. Phospholipid responses under salt- and osmotic stress

Previously, salt (NaCl) and osmotic (sorbitol, PEG, mannitol) stress have been described to rapidly trigger changes in the phospholipid profile. More precisely, PIP_2_ and PA levels were found to increase while PIP levels decreased (DeWald et al., 2001; Konig et al., 2007; Meijer et al., 2001; Meijer et al., 2017; Mishkind et al., 2009; Munnik et al., 2000; Takahashi et al., 2001; van Leeuwen et al., 2007; Zhang et al., 2018a,b).

To investigate the lipid responses in the *PLC4-KO* and -OE backgrounds, WT and mutant seedlings were prelabelled O/N with ^32^P_i_ and the next day treated for 30 min with either NaCl or sorbitol. Lipids where extracted and the levels of PIP_2_, PIP and PA measured as percentage of total ^32^P-labelled lipids (Fig. 4). In response to salt stress, PIP_2_ levels went up significantly (Fig. 4A). Less significant but still clear was the increase in PA and the small decrease in PIP (Figs. 4B, C). Interestingly, however, no significant differences between WT, *plc4* or *PLC4-OE* genotypes were observed, neither at control nor at salt stress conditions (Figs. 4A-C).

**Figure 4.**
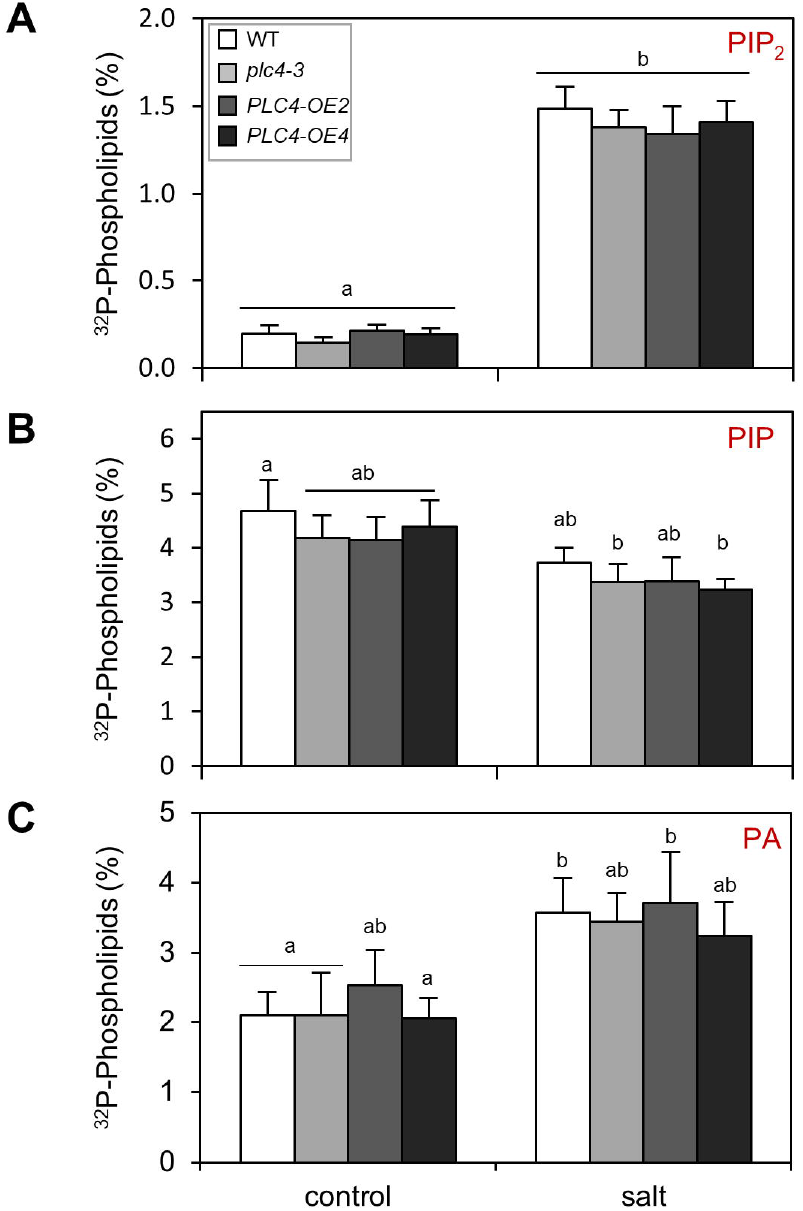
PPI- and PA levels in WT and *PLC4-KO* and -OE mutants in response to salinity stress. Three, Five-day-old seedlings, were grouped, labelled overnight with ^32^ PO_4_^3-^ and the next day treated with½ MS medium ± 300 mM NaCl for 30 min. Lipids were then extracted, separated by TLC, and quantified by phosphoimaging. ^32^P-levels of PIP_2_ (A), PIP (B) and PA (C) in WT, *plc4-3, PLC4-OE* lines #2 and #4 ± salt stress. N=7-13, from 3 to 5 different independent experiments. Error bars are SE. Samples were tested for significance using a two-way ANOVA test and post hoc TUKEY test. Significance group(s) are indicated with letters (P<0.05)

Earlier, enhanced PIP_2_ responses with sorbitol were found for Arabidopsis lines overexpressing *PLC3* and *PLC5*, but not with *PLC7*, (van Wijk et al., 2018; Zhang et al., 2018a; Zhang et al., 2018b). Here, sorbitol treatment was found to stimulate PIP_2_ and PA formation, and reduced the PIP levels, but again, no significant differences between WT and the various *PLC4* mutants were observed, as was with salt stress (Figs. 5A-C).

**Figure 5.**
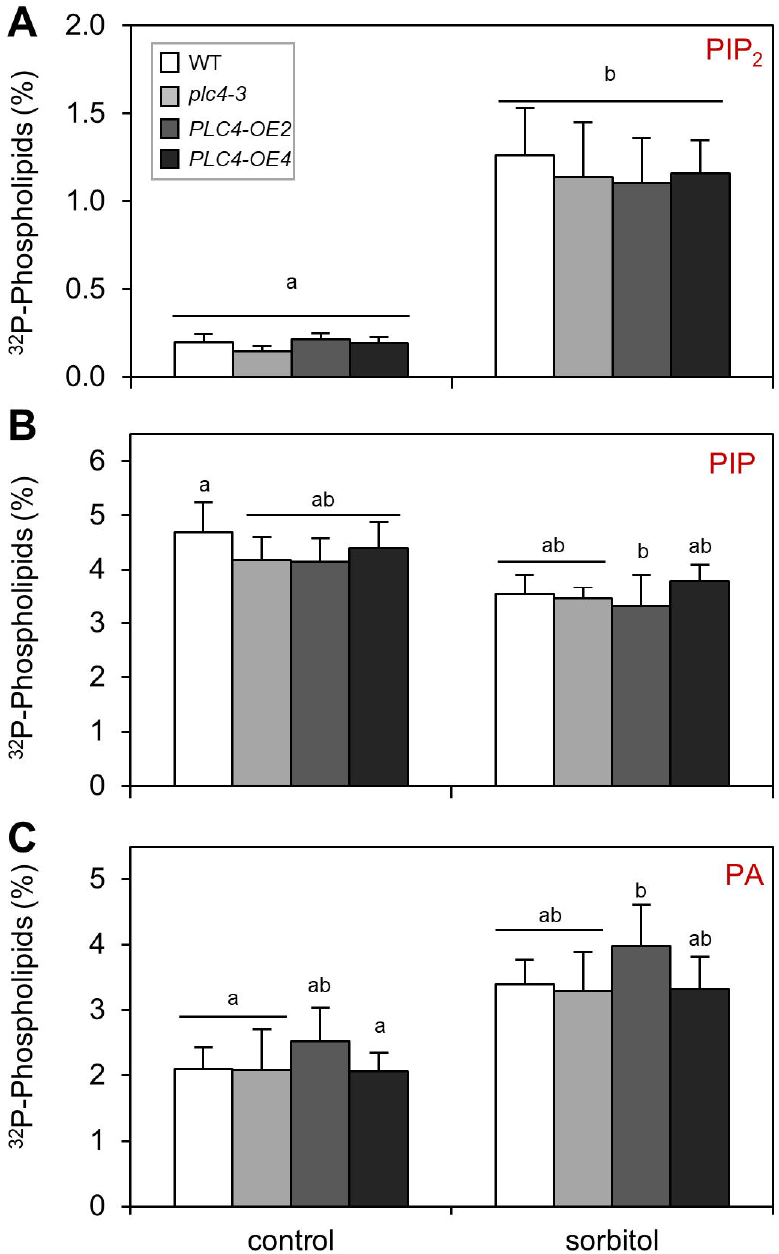
PPI- and PA levels in WT and PLC4-KO and -OE mutants in response to hyperosmotic stress. Three, Five-day-old seedlings, were grouped, labelled overnight with ^32^ PO_4_^3-^ and the next day treated with½ MS medium ± 600 mM sorbitol for 30 min. Lipids were then extracted, separated by TLC, and quantified by phosphoimaging. Data shown are the ^32^P-levels of PIP_2_ **(A)**, PIP **(B**) and PA (**C**) in WT, *plc4-3*, and *PLC4* OE lines #2 and #4 ± sorbitol. N=6-13, from 3 to 5 different independent experiments. Error bars are SE. Samples were tested for significance using a two-way ANOVA test and post hoc TUKEY test. Significance group(s) are indicated with letters (P<0.05)

In order to make sure earlier effects were not missed, lipid responses were also measured after 5 min of salt- or sorbitol stress (Fig. 6). While the PA responses became stronger, because they are faster than the PIP_2_ response, again no significant differences between WT and *PLC4* genotypes were obtained.

**Figure 6.**
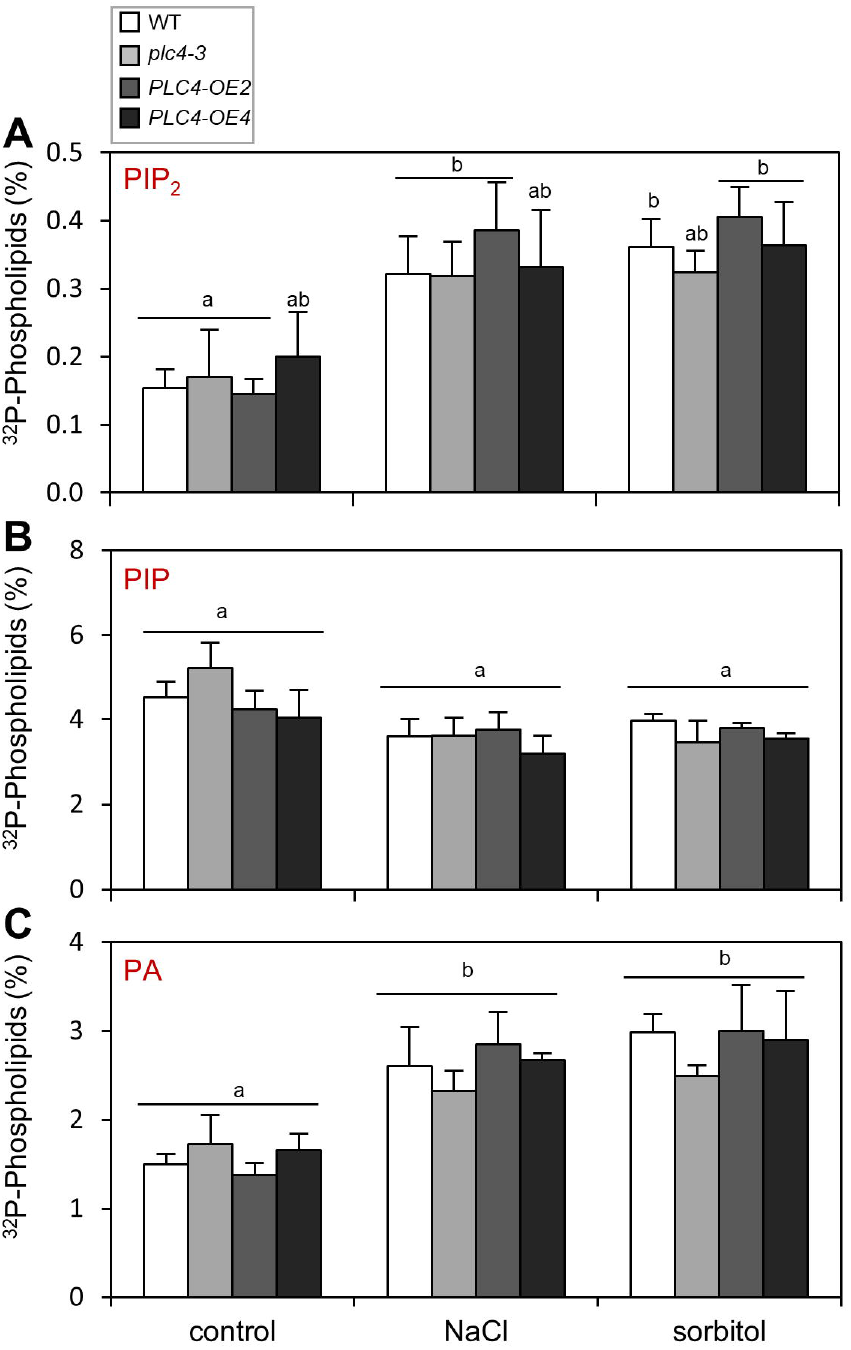
Rapid PPI- and PA responses in WT and PLC4-KO and -OE mutants in response to salt- or osmotic stress. Three, Five-day-old seedlings, were grouped, labelled overnight with ^32^ PO_4_^3-^ and the next day treated with½ MS medium ± 300 mM NaCl or 600 mM sorbitol for 5 min. Lipids were then extracted, separated by TLC, and quantified by phosphoimaging. Data shown are the ^32^P-levels of PIP_2_ (A), PIP (B) and PA (C) in WT, *plc4-3*, and *PLC4* OE lines #2 and #4 under control conditions salt- or sorbitol-stress, with N= from 5-10, from 2 to 4 different independent experiments. Error bars are SE. Samples were tested for significance using a two-way ANOVA test and post hoc TUKEY test. Significance group(s) are indicated with letters (P<0.05)

In summary, while PLC4 had been implicated in salinity stress earlier (Xia *et al*., 2017), we were unable to find significant differences in either primary root growth, or in the lipid signalling responses related to PLC, neither in salt- or osmotic stress conditions.

### 2.4. Overexpression of *PLC4* increases drought stress survival in soil

Earlier, our lab showed that ectopic overexpression of *PLC2, PLC3, PLC5, PLC7* or *PLC9* in Arabidopsis increased the survival rates when plants were exposed to drought stress (van Wijk et al., 2018; Zhang et al., 2018a; Zhang et al., 2018b; van Hooren et al., 2023). Testing this drought survival for *PLC4-OE* lines #2 and #4 grown on soil, a ∼2.5 times higher survival rate than for WT plants was obtained (Fig. 7). These results again confirm that PLC overexpression in general leads to an improved dehydration tolerance for plants and that *PLC4* make no exception.

**Figure 7.**
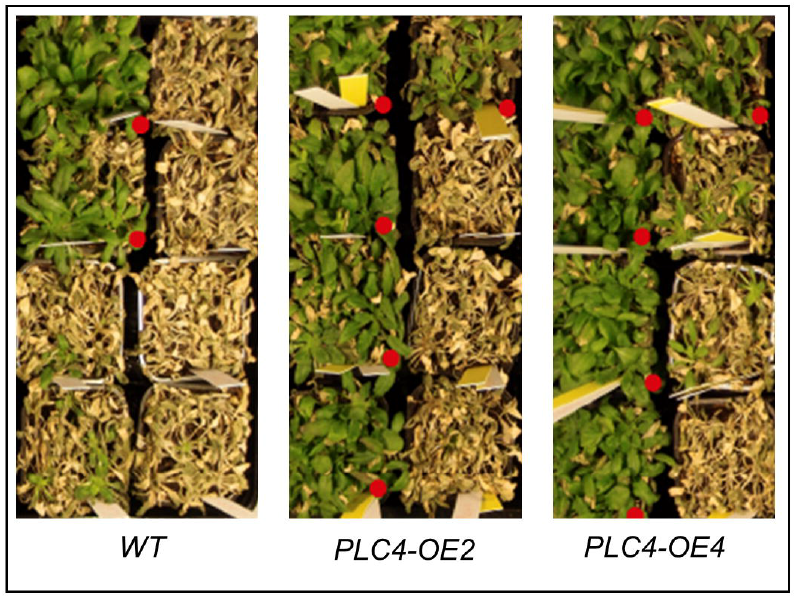
Overexpression of *PLC4* improves drought tolerance in Arabidopsis. Three weeks old plants were withheld from water for three weeks, after which they were rewatered again. Five days after rewatering they were ordered by survivors per pot and photographed. Red dots indicate surviving plants.

## 3. DISCUSSION

Overexpression of PLC has been shown to improve drought tolerance in various plant species, including Arabidopsis *PLC2, -3, -5, -7, and -9*, of which some also showed improved tolerance to salt- and osmotic stress (Deng et al., 2019; Georges et al., 2009; Tripathy et al., 2012; van Wijk et al., 2018; Wang et al., 2008; Wang et al., 2020; Zhang et al., 2018a; Zhang et al., 2018b; van Hooren et al., 2023). Recently, Xia et al. (2017) reported an opposite effect on salt tolerance for Arabidopsis *PLC4*: overexpressors being more sensitive to NaCl, while KO mutants were more tolerant. Since this observation was rather striking and unexpected, we decided to test these lines ourselves. Also, because they would make an excellent exception to the rule. However, no significant changes compared to WT were found with respect to primary root growth, nor in the lipid responses related to PLC signalling, i.e. PA, PIP and PIP_2_.

Differences between our results might be explained by the mode of overexpression. We used constitutive overexpression with the 35S promoter while Xia et al. (2017) used inducible overexpression with dexamethason. Our constitutively overexpressed-*PLC4* lines might have adapted to new equilibria (e.g. phosphoinositides, PA and other downstream processes), where the inducible lines used by Xia et al. (2017) would not have had the time to adapt to the overexpression before being exposed to the salt stress.

Another difference is that our *PLC4-OE* lines contained a C-terminal GFP fusion. Theoretically, GFP might interfere with PLC4’s function and therefore undo the OE effects. However, we could not find any GFP fluorescence for these lines under the confocal microscope, while we did find increased *PLC4* transcript levels by qPCR (Fig. 1). Moreover, increased survival rates upon drought were found for both independent *PLC4-OE* lines, indicating that PLC4 is functional.

Xia *et al*. (2017) proposed that overexpression of *PLC4* would lead to enhanced PIP_2_ hydrolysis, which would increase intracellular IP_3_- and IP_6_ concentrations and release Ca^2+^ under salt stress, so they proposed that this increase in Ca^2+^ would lead to a decrease in salt tolerance. Besides the fact that we found no change in PIP or PIP_2_ hydrolysis, we would argue that an increased Ca^2+^ response would result in an increased stress response and that this increased stress response would enable plants to survive for longer periods under severe stress conditions, as seen in our, but also in other people’s, salt- and osmotic stress experiments (Kollist et al., 2019; Kudla et al., 2018; Liu et al., 2020).

Earlier, differences in the basal level of PIP_2_ or its responses have been found for Arabidopsis *PLC-OE* lines. For example, *PLC3-OE* showed a stronger PIP_2_ responses upon sorbitol treatment (Zhang et al., 2018a), while *PLC5-OE* lines contained much lower PIP_2_ levels at control conditions, leading to relatively increased PIP_2_ responses to sorbitol as compared to WT (Zhang et al., 2018b). For *PLC7-OE* lines, no differences were found (van Wijk et al., 2018), eventhough *PLC3* and *PLC7* belong to the same clade within the *PLC* gene family (Tasma et al., 2008). *PLC4* and *PLC5* also belong to the same clade, but no difference in the basal or response levels of PIP_2_ in the *PLC4-OE* lines were observed. *PLC5-OE* showed a strong root hair phenotype (less and shorter), which turned out to be linked to the disappearance of PIP_2_ from the tip of the growing root hair (Zhang et al., 2018b). Such phenotype was not observed for *PLC4-OE* lines, nor for *PLC2, PLC3-*, *PLC7*- and *PLC9-OE* lines (van Wijk et al., 2018; Zhang et al., 2018a,b; van Hooren et al., 2023).

Besides OE of *PLC2, -3, -4, -5,-7*, and *-9* in Arabidopsis, the increase in drought tolerance has been found in various other plant systems, including maize (Wang et al., 2008), canola (Georges et al., 2009), tobacco (Tripathy et al., 2012), rice (Deng et al., 2019), wheat (Wang et al., 2020), and soybean (Chen et al., 2021a), strongly suggesting that this is a general effect of *PLC* overexpression. That plant PLCs can play an important role in stress signalling is also evident from other lines of research. Arabidopsis contains nine different *PLC* genes, which show specific and differential inductions by various biotic- and abiotic stresses (Tasma et al., 2008), providing enough redundancy to fine-tune responses and to also compensate effects of KO mutants. Aside from *plc4-KO* mutants, KO mutants of *plc2, plc3, plc7* and *plc9* and KD mutants of *plc5* have been investigated. For *plc9* mutants, reduced thermotolerance has been reported while PLC9-OE lines revealed increased tolerance (Zheng *et al*., 2012). For *plc3* and *plc5* mutants, small effects on primary- and lateral root formation were found, without getting additive effects in *plc3 plc5*-double mutants (Zhang et al., 2018a; Zhang et al., 2018b). In contrast, *plc5 plc7*-double mutants gave several additional phenotypes, including changes in stomatal movement, seed mucilage and leaf serration (van Wijk et al., 2018; Zhang et al., 2018a; Zhang et al., 2018b), while the *plc3 plc7* combination was found to be homozygous lethal (van Wijk et al., 2018). Similarly, *plc2* KO is homozygous lethal (Di Fino et al., 2017). A few ‘mutant escapes’ revealed a defect in female gametogenesis, making it sterile (Di Fino et al., 2017). PLC2 is also required for plant defence (D’Ambrosio et al., 2017) and is the only *PLC* gene of Arabidopsis that is constitutively expressed (Tasma et al., 2008; van Hooren et al., 2023). There appears to be ecotype-specific differences as well since a *plc2 KO* in the Arabidopsis Ws background is viable (Kanehara et al., 2015); all other described *plc* mutants are within the *Col- 0* background. In general, issues of lethality, or lack of strong phenotypes might be overcome by generating inducible-KO lines, to find instant disruptive phenotypes in the processes in which the particular PLC is directly involved with rather than working with a mutant who had a chance to compensate its disabilities. Since the effect of PLC could be very local, substrate and product formation may be better linked by expressing and imaging the various lipid biosensors (van Leeuwen et al., 2006; Vermeer et al., 2007; Vermeer et al., 2009; Simon et al., 2014; Vermeer et al., 2017; Li et al., 2023;). Unfortunately, there are no biosensors available for IPPs yet (de Jong and Munnik, 2021).

Many of the investigated PLCs, when constitutively overexpressed, have been associated with drought and salt, but also heat stress (Abreu et al., 2018; Chen et al., 2021b; Deng et al., 2019; Gao et al., 2014; Georges et al., 2009; Li et al., 2017; Ren et al., 2017; Tripathy et al., 2012; van Wijk et al., 2018; Wang et al., 2008; Wang et al., 2020; Zhang et al., 2018a; Zhang et al., 2018b; Zheng et al., 2012), abiotic stresses that often occur together in natural conditions. As the mutants show relatively few phenotypes, we think that the balance in phospholipid metabolism, as well as other pathways, might be shifted in these mutants so growth is not hampered.

## 4. EXPERIMENTAL

### 4.1. Arabidopsis lines

*Arabidopsis thaliana* (Col-0) was used as wild type (WT), and this ecotype is also the genetic background for all mutant lines used in this study. KO lines plc4-2 (CS876876/ SAIL_791_G05) was obtained from Nakamura’s lab (Kanehara *et al*., 2015) while plc4-3 (SALK_201150) was obtained from the SALK collection (singal.salk.edu) and was previously described by the Ren lab (Xia et al., 2017). Two lines with 35S promoter::*PLC4-GFP* i.e. *PLC4*-*OE2, PLC4-OE4* constructs were generously provided by Rodrigo Gutiérrez (Universidad Catolica de Chile) (Riveras et al., 2015).

### 4.2. Plant growth conditions

Plants were grown either on agar plates or in soil. On agar plates, plants were grown essentially as described earlier (Zhang et al., 2018b). In short, seeds were surface sterilized, stratified for two days in the dark at 4°C, and then transferred to½MS medium, supplemented with 1% Daishin agar and 1% sucrose. Plates were cultivated in growth chambers with 16-hour light/8-hour dark at 21°C. To test the effect of salt- or osmotic stress, seedlings were first grown on normal medium for four days and then transferred to plates with or without 100 mM NaCl or 200 mM sorbitol. Eight days after transfer, seedlings were scanned with an Epson Perfection V700 digital scanner and the primary root length determined using FIJI software with the SmartRoot plugin (Lobet et al., 2011; Schindelin et al., 2012). For each data point 75-99 replicates were measured over two separate experiments. Changes were verified using two-way analysis of variance (ANOVA) followed by post hoc Tukey tests.

For drought tolerance experiments, nine plants per pot (4.5×4.5×7.5cm) were grown in soil (Zaaigrond nr. 1, SIR 27010-15, JongKind BV, The Netherlands) for three weeks and watered every other day for constant water availability in the tray. After these three weeks, excess of water was removed and plants were withheld of water for another three weeks. Five days after rewatering, survivors would become green again and scored.

### 4.3. *PLC4* expression analysis

Expression levels of PLC4 in OE- and KO lines were measured using qPCR. Total RNA was extracted using Trizol reagent (Invitrogen, Carlsbad,CA). 0.5 µg of RNA isolated from six-day old seedlings was converted into cDNA using oligo-dT18 primers, dNTPs, and SuperScript III Reverse Transcriptase (Invitrogen) according to the manufacturer’s instructions. An AB 7300 Real-Time PCR system (applied Biosystems) was used for the qPCR. Relative expression levels were determined by comparing the threshold cycle values of PLC4 to the housekeeping gene, SAND (AT2G28390). Primers that were used are: PLC4_qPCR_fwd: ‘CGGAGCTCAAATGATTGC’, PLC4_qPCR_rev: ‘GTCCATTAGGACTTGCATCC’; SAND_qPCR_fwd: ‘AACTCTATGCAGCATTTGATCCACT’ and SAND_qPCR_rev: ‘TGATTGCATATCTTTATCGCCATC’. Three technical replicates and three biological replicates were used. Changes were verified using a student T-test.

### 4.4. 32 P_i_-labelling and lipid responses

Experiments were essentially performed as described earlier (Munnik and Zarza, 2013). Per sample, 3 five-day old seedlings were labelled overnight (O/N) with 5-10 µCi ^32^P-orthophosphate (^32^P_i_). The next day, seedlings were treated for 5 or 30 min in labelling buffer with or without 300 mM NaCl or 600 mM sorbitol for salt- or osmotic stress, respectively. These concentrations have similar osmolalities and are commonly used in such short-term experiments (Munnik and Vermeer, 2010). Lipids were then extracted, separated by thin layer chromatography (TLC) using an alkaline TLC solvent system that separates PIP, PIP_2_ and PA from the rest of the phospholipids (Munnik et al., 1994), quantified by phosphoimaging (Typhoon FLA 7000; GE healthcare), and expressed as a percentage of total ^32^P-labelled phospholipids. Changes were verified using two-way analysis of variance (ANOVA) followed by post hoc Tukey tests.

## ACKNOWLEDGEMENTS

We thank Rodrigo Gutiérrez (Universidad Catolica de Chile) for the *PLC4-OE* lines and Michel Haring for critical reading of the manuscript.

## DATA AVAILABILITY STATEMENT

All data supporting the findings of this study are available from the corresponding author (Teun Munnik)

## REFERENCES

Abreu, F. R. M., Dedicova, B., Vianello, R. P., Lanna, A. C., de Oliveira, J. A. V., Vieira, A. F., Morais, O. P., Mendonça, J. A., Brondani, C., 2018. Overexpression of a phospholipase (OsPLDα1) for drought tolerance in upland rice (Oryza sativa L.). Protoplasma 255, 1751–1761; https://doi.org/10.1007/s00709-018-1265-6.

Arisz, S. A., Munnik, T., 2013. Distinguishing phosphatidic acid pools from de novo synthesis, PLD, and DGK. Methods Mol Biol 1009, 55–62; https://doi.org/10.1007/978-1-62703-401-2_6.

Boretti, A., Rosa, L., 2019. Reassessing the projections of the World Water Development Report. Npj Clean Water 2; https://doi.org/10.1038/s41545-019-0039-9.

Chapman, J. M., Muhlemann, J. K., Gayomba, S. R., Muday, G. K., 2019. RBOH-Dependent ROS Synthesis and ROS Scavenging by Plant Specialized Metabolites To Modulate Plant Development and Stress Responses. Chem. Res. Toxicol. 32, 370–396; https://doi.org/10.1021/acs.chemrestox.9b00028.

Chen, L., Yang, H., Fang, Y., Guo, W., Chen, H., Zhang, X., Dai, W., Chen, S., Hao, Q., Yuan, S., Zhang, C., Huang, Y., Shan, Z., Yang, Z., Qiu, D., Liu, X., Tran, L. P., Zhou, X., Cao, D., 2021a. Overexpression of GmMYB14 improves high-density yield and drought tolerance of soybean through regulating plant architecture mediated by the brassinosteroid pathway. Plant Biotechnol. J. 19, 702–716; https://doi.org/10.1111/pbi.13496.

Chen, Z. F., Ru, J. N., Sun, G. Z., Du, Y., Chen, J., Zhou, Y. B., Chen, M., Ma, Y. Z., Xu, Z. S., Zhang, X. H., 2021b. Genomic-Wide Analysis of the PLC Family and Detection of GmPI-PLC7 Responses to Drought and Salt Stresses in Soybean. Front Plant Sci 12, 631470; https://doi.org/10.3389/fpls.2021.631470.

D’Ambrosio JM, Couto D, Fabro G, Scuffi D, Lamattina L, Munnik T, Andersson MX, Álvarez ME, Zipfel C, Laxalt AM., 2017. Phospholipase C2 Affects MAMP-Triggered Immunity by Modulating ROS Production. Plant Physiol. 175, 970–981.https://dio.org/10.1104/pp.17.00173.

de Jong, F., Munnik, T., 2021. Attracted to membranes: lipid-binding domains in plants. Plant Physiol. 185, 707–723; https://doi.org/10.1093/plphys/kiaa100.

Deng, X., Yuan, S., Cao, H., Lam, S. M., Shui, G., Hong, Y., Wang, X., 2019. Phosphatidylinositol-hydrolyzing phospholipase C4 modulates rice response to salt and drought. Plant, Cell Environ. 42, 536–548; https://doi.org/10.1111/pce.13437.

DeWald, D. B., Torabinejad, J., Jones, C. A., Shope, J. C., Cangelosi, A. R., Thompson, J. E., Prestwich, G. D., Hama, H., 2001. Rapid accumulation of phosphatidylinositol 4,5-bisphosphate and inositol 1,4,5-trisphosphate correlates with calcium mobilization in salt-stressed arabidopsis. Plant Physiol. 126, 759–769; https://doi.org/10.1104/pp.126.2.759.

Di Fino, L. M., D’Ambrosio, J. M., Tejos, R., van Wijk, R., Lamattina, L., Munnik, T., Pagnussat, G. C., Laxalt, A. M., 2017. Arabidopsis phosphatidylinositol-phospholipase C2 (PLC2) is required for female gametogenesis and embryo development. Planta 245, 717–728; https://doi.org/10.1007/s00425-016-2634-z.

Dieck, C. B., Boss, W. F., Perera, I. Y., 2012. A role for phosphoinositides in regulating plant nuclear functions. Front Plant Sci 3, 50; https://doi.org/10.3389/fpls.2012.00050.

Doumane, M., Lebecq, A., Colin, L., Fangain, A., Stevens, F. D., Bareille, J., Hamant, O., Belkhadir, Y., Munnik, T., Jaillais, Y., Caillaud, M. C., 2021. Inducible depletion of PI(4,5)P(2) by the synthetic iDePP system in Arabidopsis. Nat Plants 7, 587–597; https://doi.org/10.1038/s41477-021-00907-z.

Flores, S., Smart, C. C., 2000. Abscisic acid-induced changes in inositol metabolism in Spirodela polyrrhiza. Planta 211, 823–832; https://doi.org/10.1007/s004250000348.

Gamalero, E., Glick, B. R., 2022. Recent Advances in Bacterial Amelioration of Plant Drought and Salt Stress. Biology (Basel) 11, 437; https://doi.org/10.3390/biology11030437.

Gao, K., Liu, Y. L., Li, B., Zhou, R. G., Sun, D. Y., Zheng, S. Z., 2014. Arabidopsis thaliana phosphoinositide-specific phospholipase C isoform 3 (AtPLC3) and AtPLC9 have an additive effect on thermotolerance. Plant Cell Physiol 55, 1873–1883; https://doi.org/10.1093/pcp/pcu116.

Georges, F., Das, S., Ray, H., Bock, C., Nokhrina, K., Kolla, V. A., Keller, W., 2009. Over-expression of Brassica napus phosphatidylinositol-phospholipase C2 in canola induces significant changes in gene expression and phytohormone distribution patterns, enhances drought tolerance and promotes early flowering and maturation. Plant, Cell Environ. 32, 1664–1681; https://doi.org/10.1111/j.1365-3040.2009.02027.x.

Golldack, D., Li, C., Mohan, H., Probst, N., 2014. Tolerance to drought and salt stress in plants: Unraveling the signaling networks. Front Plant Sci 5, 151; https://doi.org/10.3389/fpls.2014.00151.

Guo, T., Chen, H. C., Lu, Z. Q., Diao, M., Chen, K., Dong, N. Q., Shan, J. X., Ye, W. W., Huang, S., Lin, H. X., 2020. A SAC Phosphoinositide Phosphatase Controls Rice Development via Hydrolyzing PI4P and PI(4,5)P2. Plant Physiol. 182, 1346–1358; https://doi.org/10.1104/pp.19.01131.

Han, X., Yang, Y., 2021. Phospholipids in Salt Stress Response. Plants (Basel) 10, 2204; https://doi.org/10.3390/plants10102204.

Hao, Z., Ma, S., Liang, L., Feng, T., Xiong, M., Lian, S., Zhu, J., Chen, Y., Meng, L., Li, M., 2022. Candidate Genes and Pathways in Rice Co-Responding to Drought and Salt Identified by gcHap Network. Int. J. Mol. Sci. 23, 4016; https://doi.org/10.3390/ijms23074016.

Hou, Q., Ufer, G., Bartels, D., 2016. Lipid signalling in plant responses to abiotic stress. Plant Cell Environ. 39, 1029–1048; https://doi.org/10.1111/pce.12666.

Hunt, L., Mills, L. N., Pical, C., Leckie, C. P., Aitken, F. L., Kopka, J., Mueller-Roeber, B., McAinsh, M. R., Hetherington, A. M., Gray, J. E., 2003. Phospholipase C is required for the control of stomatal aperture by ABA. Plant J. 34, 47–55; https://doi.org/10.1046/j.1365-313x.2003.01698.x.

Ischebeck, T., Seiler, S., Heilmann, I., 2010. At the poles across kingdoms: phosphoinositides and polar tip growth. Protoplasma 240, 13–31; https://doi.org/10.1007/s00709-009-0093-0.

Ischebeck, T., Werner, S., Krishnamoorthy, P., Lerche, J., Meijon, M., Stenzel, I., Lofke, C., Wiessner, T., Im, Y. J., Perera, I. Y., Iven, T., Feussner, I., Busch, W., Boss, W. F., Teichmann, T., Hause, B., Persson, S., Heilmann, I., 2013. Phosphatidylinositol 4,5-bisphosphate influences PIN polarization by controlling clathrin-mediated membrane trafficking in Arabidopsis. Plant Cell 25, 4894–4911; https://doi.org/10.1105/tpc.113.116582.

Kadota, Y., Shirasu, K., Zipfel, C., 2015. Regulation of the NADPH Oxidase RBOHD During Plant Immunity. Plant Cell Physiol 56, 1472–1480; https://doi.org/10.1093/pcp/pcv063.

Kanehara, K., Yu, C.-Y., Cho, Y., Cheong, W.-F., Torta, F., Shui, G., Wenk, M. R., Nakamura, Y., 2015. Arabidopsis AtPLC2 is a primary phosphoinositide-specific phospholipase C in phosphoinositide metabolism and the endoplasmic reticulum stress response. PLoS Genet 11, e1005511; https://doi.org/10.1371/journal.pgen.1005511.

Katagiri, T., Takahashi, S., Shinozaki, K., 2001. Involvement of a novel Arabidopsis phospholipase D, AtPLDdelta, in dehydration-inducible accumulation of phosphatidic acid in stress signalling. Plant J. 26, 595–605; https://doi.org/10.1046/j.1365-313x.2001.01060.x.

Kato, M., Tsuge, T., Maeshima, M., Aoyama, T., 2019. Arabidopsis PCaP2 modulates the phosphatidylinositol 4,5-bisphosphate signal on the plasma membrane and attenuates root hair elongation. Plant J. 99, 610–625; https://doi.org/10.1111/tpj.14226.

Kollist, H., Zandalinas, S. I., Sengupta, S., Nuhkat, M., Kangasjarvi, J., Mittler, R., 2019. Rapid Responses to Abiotic Stress: Priming the Landscape for the Signal Transduction Network. Trends Plant Sci. 24, 25–37; https://doi.org/10.1016/j.tplants.2018.10.003.

Konig, S., Mosblech, A., Heilmann, I., 2007. Stress-inducible and constitutive phosphoinositide pools have distinctive fatty acid patterns in Arabidopsis thaliana. FASEB J. 21, 1958–1967; https://doi.org/10.1096/fj.06-7887com.

Kudla, J., Becker, D., Grill, E., Hedrich, R., Hippler, M., Kummer, U., Parniske, M., Romeis, T., Schumacher, K., 2018. Advances and current challenges in calcium signaling. New Phytol 218, 414–431; https://doi.org/10.1111/nph.14966.

Kuroda, R., Kato, M., Tsuge, T., Aoyama, T., 2021. Arabidopsis phosphatidylinositol 4-phosphate 5-kinase genes PIP5K7, PIP5K8, and PIP5K9 are redundantly involved in root growth adaptation to osmotic stress. Plant J. 106, 913–927; https://doi.org/10.1111/tpj.15207.

Laha, D., Parvin, N., Dynowski, M., Johnen, P., Mao, H., Bitters, S. T., Zheng, N., Schaaf, G., 2016. Inositol Polyphosphate Binding Specificity of the Jasmonate Receptor Complex. Plant Physiol. 171, 2364–2370; https://doi.org/10.1104/pp.16.00694.

Lebecq, A., Doumane, M., Fangain, A., Bayle, V., Leong, J. X., Rozier, F., Marques-Bueno, M. D., Armengot, L., Boisseau, R., Simon, M. L., Franz-Wachtel, M., Macek, B., Ustun, S., Jaillais, Y., Caillaud, M. C., 2022. The Arabidopsis SAC9 enzyme is enriched in a cortical population of early endosomes and restricts PI(4,5)P2 at the plasma membrane. Elife 11; https://doi.org/10.7554/eLife.73837.

Lemtiri-Chlieh, F., MacRobbie, E. A., Brearley, C. A., 2000. Inositol hexakisphosphate is a physiological signal regulating the K+-inward rectifying conductance in guard cells. Proc Natl Acad Sci U S A 97, 8687–8692; https://doi.org/10.1073/pnas.140217497.

Lemtiri-Chlieh, F., MacRobbie, E. A., Webb, A. A., Manison, N. F., Brownlee, C., Skepper, J. N., Chen, J., Prestwich, G. D., Brearley, C. A., 2003. Inositol hexakisphosphate mobilizes an endomembrane store of calcium in guard cells. Proceedings of the National Academy of Sciences 100, 10091–10095; https://doi.org/10.1073/pnas.1133289100.

Li, L., Wang, F., Yan, P., Jing, W., Zhang, C., Kudla, J., Zhang, W., 2017. A phosphoinositide-specific phospholipase C pathway elicits stress-induced Ca2+ signals and confers salt tolerance to rice. New Phytol. 214, 1172–1187; https://doi.org/10.1111/nph.14426.

Li, T., Xiao, X., Liu, Q., Li, W., Li, L., Zhang, W., Munnik, T., Wang, X., Zhang Q (2023). Dynamic responses of PA to environmental stimuli imaged by a genetically encoded mobilizable fluorescent sensor. Plant Commun. 4: 100500; https://doi.org/10.1016/j.xplc.2022.100500

Liu, J., Lenzoni, G., Knight, M. R., 2020. Design Principle for Decoding Calcium Signals to Generate Specific Gene Expression Via Transcription. Plant Physiol. 182, 1743–1761; https://doi.org/10.1104/pp.19.01003.

Lobet, G., Pagès, L., Draye, X., 2011. A novel image-analysis toolbox enabling quantitative analysis of root system architecture. Plant Physiol. 157, 29–39; https://doi.org/10.1104/pp.111.179895.

Marusig, D., Tombesi, S., 2020. Abscisic Acid Mediates Drought and Salt Stress Responses in Vitis vinifera-A Review. Int. J. Mol. Sci. 21, 8648; https://doi.org/10.3390/ijms21228648.

Mei, Y., Jia, W. J., Chu, Y. J., Xue, H. W., 2012. Arabidopsis phosphatidylinositol monophosphate 5-kinase 2 is involved in root gravitropism through regulation of polar auxin transport by affecting the cycling of PIN proteins. Cell Res. 22, 581–597; https://doi.org/10.1038/cr.2011.150.

Meijer, H. J., Arisz, S. A., Van Himbergen, J. A., Musgrave, A., Munnik, T., 2001. Hyperosmotic stress rapidly generates lyso-phosphatidic acid in Chlamydomonas. Plant J. 25, 541–548; https://doi.org/10.1046/j.1365-313x.2001.00990.x.

Meijer, H. J., van Himbergen, J. A., Musgrave, A., Munnik, T., 2017. Acclimation to salt modifies the activation of several osmotic stress-activated lipid signalling pathways in Chlamydomonas. Phytochemistry 135, 64–72; https://doi.org/10.1016/j.phytochem.2016.12.014.

Mills, L. N., Hunt, L., Leckie, C. P., Aitken, F. L., Wentworth, M., McAinsh, M. R., Gray, J. E., Hetherington, A. M., 2004. The effects of manipulating phospholipase C on guard cell ABA-signalling. J Exp Bot 55, 199–204; https://doi.org/10.1093/jxb/erh027.

Mishkind, M., Vermeer, J. E., Darwish, E., Munnik, T., 2009. Heat stress activates phospholipase D and triggers PIP accumulation at the plasma membrane and nucleus. Plant J. 60, 10–21; https://doi.org/10.1111/j.1365-313X.2009.03933.x.

Munnik, T., 2014. PI-PLC: phosphoinositide-phospholipase C in plant signaling. Phospholipases in Plant Signaling. Springer, pp. 27–54

Munnik, T., Irvine, R. F., Musgrave, A., 1994. Rapid turnover of phosphatidylinositol 3-phosphate in the green alga Chlamydomonas eugametos: signs of a phosphatidylinositide 3-kinase signalling pathway in lower plants? Biochem J 298 (Pt2), 269–273; https://doi.org/10.1042/bj2980269.

Munnik, T., Meijer, H. J., Ter Riet, B., Hirt, H., Frank, W., Bartels, D., Musgrave, A., 2000. Hyperosmotic stress stimulates phospholipase D activity and elevates the levels of phosphatidic acid and diacylglycerol pyrophosphate. Plant J. 22, 147–154; https://doi.org/10.1046/j.1365-313x.2000.00725.x.

Munnik, T., Testerink, C., 2009. Plant phospholipid signaling: “in a nutshell”. J Lipid Res 50 Suppl, S260–265; https://doi.org/10.1194/jlr.R800098-JLR200.

Munnik, T., Vermeer, J. E., 2010. Osmotic stress-induced phosphoinositide and inositol phosphate signalling in plants. Plant Cell Environ. 33, 655–669; https://doi.org/10.1111/j.1365-3040.2009.02097.x.

Munnik, T., Zarza, X., 2013. Analyzing plant signaling phospholipids through 32Pi-labeling and TLC. Methods Mol Biol 1009, 3–15; https://doi.org/10.1007/978-1-62703-401-2_1.

Naramoto, S., Sawa, S., Koizumi, K., Uemura, T., Ueda, T., Friml, J., Nakano, A., Fukuda, H., 2009. Phosphoinositide-dependent regulation of VAN3 ARF-GAP localization and activity essential for vascular tissue continuity in plants. Development 136, 1529–1538; https://doi.org/10.1242/dev.030098.

Noack, L. C., Jaillais, Y., 2017. Precision targeting by phosphoinositides: how PIs direct endomembrane trafficking in plants. Curr. Opin. Plant Biol. 40, 22–33; https://doi.org/10.1016/j.pbi.2017.06.017.

Noack, L. C., Jaillais, Y., 2020. Functions of Anionic Lipids in Plants. Annu. Rev. Plant Biol. 71, 71–102; https://doi.org/10.1146/annurev-arplant-081519-035910.

Pleskot, R., Li, J., Zarsky, V., Potocky, M., Staiger, C. J., 2013. Regulation of cytoskeletal dynamics by phospholipase D and phosphatidic acid. Trends Plant Sci. 18, 496–504; https://doi.org/10.1016/j.tplants.2013.04.005.

Pokotylo, I., Kolesnikov, Y., Kravets, V., Zachowski, A., Ruelland, E., 2014. Plant phosphoinositide-dependent phospholipases C: variations around a canonical theme. Biochimie 96, 144–157; https://doi.org/10.1016/j.biochi.2013.07.004.

Pokotylo, I., Kravets, V., Martinec, J., Ruelland, E., 2018. The phosphatidic acid paradox: Too many actions for one molecule class? Lessons from plants. Prog Lipid Res 71, 43–53; https://doi.org/10.1016/j.plipres.2018.05.003.

Qiu, Q. S., Guo, Y., Quintero, F. J., Pardo, J. M., Schumaker, K. S., Zhu, J. K., 2004. Regulation of vacuolar Na+/H+ exchange in Arabidopsis thaliana by the salt-overly-sensitive (SOS) pathway. J Biol Chem 279, 207–215; https://doi.org/10.1074/jbc.M307982200.

Ren, H., Gao, K., Liu, Y., Sun, D., Zheng, S., 2017. The role of AtPLC3 and AtPLC9 in thermotolerance in Arabidopsis. Plant Signal Behav 12, e1162368; https://doi.org/10.1080/15592324.2016.1162368.

Riveras, E., Alvarez, J. M., Vidal, E. A., Oses, C., Vega, A., Gutierrez, R. A., 2015. The Calcium Ion Is a Second Messenger in the Nitrate Signaling Pathway of Arabidopsis. Plant Physiol. 169, 1397–1404; https://doi.org/10.1104/pp.15.00961.

Schindelin, J., Arganda-Carreras, I., Frise, E., Kaynig, V., Longair, M., Pietzsch, T., Preibisch, S., Rueden, C., Saalfeld, S., Schmid, B., 2012. Fiji: an open-source platform for biological-image analysis. Nat. Methods 9, 676–682; https://doi.org/10.1038/nmeth.2019.

Shen, L., Zhuang, B., Wu, Q., Zhang, H., Nie, J., Jing, W., Yang, L., Zhang, W., 2019. Phosphatidic acid promotes the activation and plasma membrane localization of MKK7 and MKK9 in response to salt stress. Plant Sci. 287, 110190; https://doi.org/10.1016/j.plantsci.2019.110190.

Simon, M.L., Platre, M.P., Assil, S., van Wijk, R., Chen, W.Y., Chory, J., Dreux, M., Munnik, T., Jaillais, Y. (2014) A multi-colour/multi-affinity marker set to visualize phosphoinositide dynamics in Arabidopsis. Plant J. 77: 322–337; https://doi.org/10.1111/tpj.12358.

Song, L., Wang, Y., Guo, Z., Lam, S. M., Shui, G., Cheng, Y., 2021. NCP2/RHD4/SAC7, SAC6 and SAC8 phosphoinositide phosphatases are required for PtdIns4P and PtdIns(4,5)P2 homeostasis and Arabidopsis development. New Phytol 231, 713–725; https://doi.org/10.1111/nph.17402.

Synek, L., Pleskot, R., Sekeres, J., Serrano, N., Vukasinovic, N., Ortmannova, J., Klejchova, M., Pejchar, P., Batystova, K., Gutkowska, M., Jankova-Drdova, E., Markovic, V., Pecenkova, T., Santrucek, J., Zarsky, V., Potocky, M., 2021. Plasma membrane phospholipid signature recruits the plant exocyst complex via the EXO70A1 subunit. Proc Natl Acad Sci U S A 118; https://doi.org/10.1073/pnas.2105287118.

Takahashi, S., Katagiri, T., Hirayama, T., Yamaguchi-Shinozaki, K., Shinozaki, K., 2001. Hyperosmotic stress induces a rapid and transient increase in inositol 1,4,5-trisphosphate independent of abscisic acid in Arabidopsis cell culture. Plant Cell Physiol 42, 214–222; https://doi.org/10.1093/pcp/pce028.

Tasma, I. M., Brendel, V., Whitham, S. A., Bhattacharyya, M. K., 2008. Expression and evolution of the phosphoinositide-specific phospholipase C gene family in Arabidopsis thaliana. Plant Physiol. Biochem. 46, 627–637; https://doi.org/10.1016/j.plaphy.2008.04.015.

Testerink, C., Munnik, T., 2011. Molecular, cellular, and physiological responses to phosphatidic acid formation in plants. J Exp Bot 62, 2349–2361; https://doi.org/10.1093/jxb/err079.

Tripathy, M. K., Tyagi, W., Goswami, M., Kaul, T., Singla-Pareek, S. L., Deswal, R., Reddy, M. K., Sopory, S. K., 2012. Characterization and functional validation of tobacco PLC delta for abiotic stress tolerance. Plant Mol. Biol. Rep. 30, 488–497; https://doi.org/10.1007/s11105-011-0360-z.

Ufer, G., Gertzmann, A., Gasulla, F., Rohrig, H., Bartels, D., 2017. Identification and characterization of the phosphatidic acid-binding A. thaliana phosphoprotein PLDrp1 that is regulated by PLDalpha1 in a stress-dependent manner. Plant J. 92, 276–290; https://doi.org/10.1111/tpj.13651.

Uraji, M., Katagiri, T., Okuma, E., Ye, W., Hossain, M. A., Masuda, C., Miura, A., Nakamura, Y., Mori, I. C., Shinozaki, K., Murata, Y., 2012. Cooperative function of PLDdelta and PLDalpha1 in abscisic acid-induced stomatal closure in Arabidopsis. Plant Physiol. 159, 450–460; https://doi.org/10.1104/pp.112.195578.

van Hooren, M., van Wijk, R., Vaseva, I.I.j., van der Straeten, D., Haring, M.A., Munnik, T. 2023. Ectopic expression of distinct PLC genes identifies ‘Compactness’ as novel architectural shoot strategy to cope with drought stress. Plant Cell Physiol. Submitted May 2023.

van Leeuwen, W., Vermeer, J. E., Gadella, T. W., Jr., Munnik, T., 2007. Visualization of phosphatidylinositol 4,5-bisphosphate in the plasma membrane of suspension-cultured tobacco BY-2 cells and whole Arabidopsis seedlings. Plant J. 52, 1014–1026; https://doi.org/10.1111/j.1365-313X.2007.03292.x.

van Wijk, R., Zhang, Q., Zarza, X., Lamers, M., Marquez, F. R., Guardia, A., Scuffi, D., Garcia-Mata, C., Ligterink, W., Haring, M. A., Laxalt, A. M., Munnik, T., 2018. Role for Arabidopsis PLC7 in Stomatal Movement, Seed Mucilage Attachment, and Leaf Serration. Front Plant Sci 9, 1721; https://doi.org/10.3389/fpls.2018.01721.

Verma, S., Negi, N. P., Pareek, S., Mudgal, G., Kumar, D., 2022. Auxin response factors in plant adaptation to drought and salinity stress. Physiol Plant 174, e13714; https://doi.org/10.1111/ppl.13714.

Vermeer JE, van Leeuwen W, Tobeña-Santamaria R, Laxalt AM, Jones DR, Divecha N, Gadella TW Jr, Munnik T., 2006. Visualization of PtdIns3P dynamics in living plant cells. Plant J. 47, 687–700. https://doi.org/10.1111/j.1365-313X.2006.02830.x

Vermeer, J.E.M., Thole, J.M., Goedhart, J., Nielsen, E., Munnik, T. and Gadella Jr, T.W.J., 2009. Imaging phosphatidylinositol 4-phosphate dynamics in living plant cells. The Plant Journal, 57, 356–37;. https://doi.org/10.1111/j.1365-313X.2008.03679.x

Vermeer JEM, van Wijk R, Goedhart J, Geldner N, Chory J, Gadella TWJ Jr, Munnik T., 2017. In Vivo Imaging of Diacylglycerol at the Cytoplasmic Leaflet of Plant Membranes. Plant Cell Physiol. 58, 1196–1207. https://doi.org/10.1093/pcp/pcx012.

Verslues, P.E, Bailey-Serres, J., Brodersen, C., Buckley, T.N., Conti, L., Christmann, A., Dinneny, J.R., Grill, E., Hayes, S., Heckman, R.W., Hsu, P.K,. Juenger, T.E., Mas, P., Munnik, T., Nelissen, H., Sack, L., Schroeder, J.I., Testerink, C., Tyerman, S.D., Umezawa, T., Wigge, P.A. (2023) Burning questions for a warming and changing world: 15 unknowns in plant abiotic stress. Plant Cell 35: 67–108; https://doi.org/10.1093/plcell/koac263

Wang, C.-R., Yang, A.-F., Yue, G.-D., Gao, Q., Yin, H.-Y., Zhang, J.-R., 2008. Enhanced expression of phospholipase C 1 (ZmPLC1) improves drought tolerance in transgenic maize. Planta 227, 1127–1140; https://doi.org/10.1007/s00425-007-0686-9.

Wang, P., Shen, L., Guo, J., Jing, W., Qu, Y., Li, W., Bi, R., Xuan, W., Zhang, Q., Zhang, W., 2019. Phosphatidic Acid Directly Regulates PINOID-Dependent Phosphorylation and Activation of the PIN-FORMED2 Auxin Efflux Transporter in Response to Salt Stress. Plant Cell 31, 250–271; https://doi.org/10.1105/tpc.18.00528.

Wang, X., Liu, Y., Li, Z., Gao, X., Dong, J., Zhang, J., Zhang, L., Thomashow, L. S., Weller, D. M., Yang, M., 2020. Genome-Wide Identification and Expression Profile Analysis of the Phospholipase C Gene Family in Wheat (Triticum aestivum L.) Plants (Basel) 9, 885; https://doi.org/10.3390/plants9070885.

Wu, C., Tan, L., van Hooren, M., Tan, X., Liu, F., Li, Y., Zhao, Y., Li, B., Rui, Q., Munnik, T., Bao, Y., 2017. Arabidopsis EXO70A1 recruits Patellin3 to the cell membrane independent of its role as an exocyst subunit. J Integr Plant Biol 59, 851–865; https://doi.org/10.1111/jipb.12578.

Xia, K., Wang, B., Zhang, J., Li, Y., Yang, H., Ren, D., 2017. Arabidopsis phosphoinositide-specific phospholipase C 4 negatively regulates seedling salt tolerance. Plant Cell Environ. 40, 1317–1331; https://doi.org/10.1111/pce.12918.

Xing, J., Zhang, L., Duan, Z., Lin, J., 2021. Coordination of Phospholipid-Based Signaling and Membrane Trafficking in Plant Immunity. Trends Plant Sci. 26, 407–420; https://doi.org/10.1016/j.tplants.2020.11.010.

Yao, H. Y., Xue, H. W., 2018. Phosphatidic acid plays key roles regulating plant development and stress responses. J. Integr. Plant Biol. 60, 851–863; https://doi.org/10.1111/jipb.12655.

Yperman, K., Wang, J., Eeckhout, D., Winkler, J., Vu, L. D., Vandorpe, M., Grones, P., Mylle, E., Kraus, M., Merceron, R., Nolf, J., Mor, E., De Bruyn, P., Loris, R., Potocky, M., Savvides, S. N., De Rybel, B., De Jaeger, G., Van Damme, D., Pleskot, R., 2021. Molecular architecture of the endocytic TPLATE complex. Sci Adv 7, eabe7999; https://doi.org/10.1126/sciadv.abe7999.

Yu, L., Nie, J., Cao, C., Jin, Y., Yan, M., Wang, F., Liu, J., Xiao, Y., Liang, Y., Zhang, W., 2010. Phosphatidic acid mediates salt stress response by regulation of MPK6 in Arabidopsis thaliana. New Phytol 188, 762–773; https://doi.org/10.1111/j.1469-8137.2010.03422.x.

Zhang, Q., van Wijk, R., Shahbaz, M., Roels, W., Schooten, B. V., Vermeer, J. E. M., Zarza, X., Guardia, A., Scuffi, D., Garcia-Mata, C., Laha, D., Williams, P., Willems, L. A. J., Ligterink, W., Hoffmann-Benning, S., Gillaspy, G., Schaaf, G., Haring, M. A., Laxalt, A. M., Munnik, T., 2018a. Arabidopsis Phospholipase C3 is Involved in Lateral Root Initiation and ABA Responses in Seed Germination and Stomatal Closure. Plant Cell Physiol 59, 469–486; https://doi.org/10.1093/pcp/pcx194.

Zhang, Q., van Wijk, R., Zarza, X., Shahbaz, M., van Hooren, M., Guardia, A., Scuffi, D., Garcia-Mata, C., Van den Ende, W., Hoffmann-Benning, S., Haring, M. A., Laxalt, A. M., Munnik, T., 2018b. Knock-Down of Arabidopsis PLC5 Reduces Primary Root Growth and Secondary Root Formation While Overexpression Improves Drought Tolerance and Causes Stunted Root Hair Growth. Plant Cell Physiol 59, 2004–2019; https://doi.org/10.1093/pcp/pcy120.

Zhang, Y., Zhu, H., Zhang, Q., Li, M., Yan, M., Wang, R., Wang, L., Welti, R., Zhang, W., Wang, X., 2009. Phospholipase Dα1 and phosphatidic acid regulate NADPH oxidase activity and production of reactive oxygen species in ABA-mediated stomatal closure in Arabidopsis. The Plant Cell 21, 2357–2377; https://doi.org/10.1105/tpc.108.062992.

Zhao, Y., Yan, A., Feijo, J. A., Furutani, M., Takenawa, T., Hwang, I., Fu, Y., Yang, Z., 2010. Phosphoinositides regulate clathrin-dependent endocytosis at the tip of pollen tubes in Arabidopsis and tobacco. Plant Cell 22, 4031–4044; https://doi.org/10.1105/tpc.110.076760.

Zheng, S. Z., Liu, Y. L., Li, B., Shang, Z. L., Zhou, R. G., Sun, D. Y., 2012. Phosphoinositide-specific phospholipase C9 is involved in the thermotolerance of Arabidopsis. Plant J. 69, 689–700; https://doi.org/10.1111/j.1365-313X.2011.04823.x.

Zonia, L. & Munnik, T., 2004. Osmotically Induced Cell Swelling versus Cell Shrinking Elicits Specific Changes in Phospholipid Signals in Tobacco Pollen Tubes. Plant physiol. 134. https://doi.org/813-23; 10.1104/pp.103.029454.

